# Morphological analysis of human and mouse dendritic spines reveals a morphological continuum and differences across ages and species

**DOI:** 10.1101/2022.01.24.477510

**Authors:** Netanel Ofer, Ruth Benavides-Piccione, Javier DeFelipe, Rafael Yuste

**Affiliations:** Neurotechnology Center, Dept. Biological Sciences, Columbia University, New York, NY 10027; Laboratorio Cajal de Circuitos Corticales, Centro de Tecnología Biomédica, Universidad Politécnica de Madrid, Pozuelo de Alarcón, 28223 Madrid, Spain; Instituto Cajal, Consejo Superior de Investigaciones Científicas, 28002 Madrid, Spain

## Abstract

Dendritic spines have diverse morphologies, with a wide range of head and neck sizes, and these morphological differences likely generate different synaptic and functional properties. To explore how this morphological diversity differs across species we analyzed 3D confocal reconstructions of ~8,000 human spines and ~1,700 mouse spines, labeled by intracellular injections in fixed tissue. Using unsupervised machine-learning algorithms, we computationally separated spine heads and necks and systematically measured morphological features of spines in apical and basal dendrites from cortical pyramidal cells. Human spines had unimodal distributions of parameters, without any evidence of morphological subtypes. Their spine necks were longer and thinner in apical than in basal spines, and spine head volumes of an 85-years-old individual were larger than those of a 40-years-old individual. Human spines overall had longer and thicker necks and bigger head volumes than mouse spines. Our results indicate that human spines form part of a morphological continuum, are larger and longer than those of mice, and become larger with increasing adult age. These morphological differences in spines across species could generate functional differences in biochemical and electrical spine compartmentalization, or in synaptic properties, across species and ages.

## INTRODUCTION

Dendritic spines, first described by Cajal (1888, 1904) and are considered key elements in learning, memory, and cognition (Yuste, 2010, 2015). Spines are sites of most excitatory synapses in many brain areas, and practically all spines receive at least one excitatory synapse (Arellano et al., 2007; Harris and Weinberg, 2012; DeFelipe, 2015). Thus, spine numbers and shape likely influence cortical functions. Differences in spine density and size between cortical areas and species exist (Elston et al., 2001, 2011; Jacobs et al., 2001, 2002; Ballesteros-Yáñez et al., 2010). Spines in human cortex have higher density, head size, and neck length than those of mice (Benavides-Piccione et al., 2002; Ballesteros-Yáñez et al., 2010). However, mouse prelimbic cortex has similar spine density as human (Ballesteros-Yáñez et al., 2010). In human neurons, spines on apical dendrites are denser, larger, and longer than those on basal dendrites (Benavides-Piccione et al., 2013). Also, age-dependent functional changes in human cortex could be mediated by age-related spine loss (Rakic et al., 1994; Jacobs et al., 1997; Hof and Morrison, 2004; Petanjek et al., 2008; Benavides-Piccione et al., 2013; Dickstein et al., 2013). Indeed, small, short spines from basal dendrites and long spines from apical dendrites are lost with age (Benavides-Piccione et al., 2013).

Light microscopy (LM) has been traditionally used to reconstruct spine structures, enabling the dynamic tracking of spine morphologies in living neurons (Fischer et al., 1998; Dunaevsky et al., 1999; Tønnesen et al., 2014; Loewenstein et al., 2015; Bokota et al., 2016; Dickstein et al., 2016; Kashiwagi et al., 2019), revealing that spines are dynamic, changing size and shape over timescales of seconds, due to actin-based motility (Fischer et al., 1998; Dunaevsky et al., 1999; Hering and Sheng, 2001). However, previous LM studies were based on manual measurements, as automatic detecting, segmenting, and measuring spines from LM images is challenging (Levet et al., 2020; Okabe, 2020). Fluorescence signals from spines are often weak, making it difficult to identify clear spine borders. Also, the spine neck is at times invisible due to its small length and diameter, often below the optical resolution limit. Thus, separating the head and neck is particularly challenging. Electron microscopy (EM) solves these issues, and enables high-resolution nanometer-scale 3D morphological analysis of dendritic spines (Ofer et al., 2021) and the reconstruction of spines located directly above or below the dendritic shaft (Parajuli and Koike, 2021). But, in spite of recent advances in large-scale automated serial EM, it still requires significant resources and time, even to reconstruct small tissue volumes. In contrast, confocal microscopy allows the visualization of thousands of spines with high signal to noise (Benavides-Piccione et al., 2013). Thus, automatic methods to measure spines structure in LM databases could enable the systematic analysis of spine morphologies in living tissue.

To develop an automatic analysis pipeline for LM images of spines, we applied unsupervised machine-learning algorithms to computationally separate and measure spine head and neck from LM images of a dataset of 3D reconstructed human and mouse dendritic spines. These tools build upon previous spine computational repairing method (Luengo-Sanchez et al., 2018) and algorithms for spine reconstructions from EM (Ofer et al., 2021). Using these computational methods, we systematically examined morphological variables of apical and basal dendritic spines from pyramidal neurons from cingulate cortex of two human individuals of different ages, and compared those measurements with those of pyramidal neurons from mouse somatosensory cortex. Our results reveal a continuum of spine morphologies, without any clear subtypes, and confirm significant differences in spine structures between species, with human spines being systematically longer and larger. These morphological differences imply the existence functional differences in synaptic and spine function across species.

## METHODS

### Human spines dataset

We used a database of 7,917 3D-reconstructed spines (Figure 1; published in Benavides-Piccione et al., 2013), from intracellularly injected apical and basal dendrites from layer 3 pyramidal neurons in the cingulate cortex of two human males aged 40 and 85 obtained at autopsy. Briefly, brains were immersed (2–3 h post-mortem), in cold 4% paraformaldehyde in 0.1 M phosphate buffer, pH 7.4 (PB) and sectioned into 1.5-cm-thick coronal slices. Vibratome sections (250 *μm*) from the anterior cingular gyri (corresponding to Brodmann’s area 24; Garey, 1999) were obtained with a vibratome and labelled with 4,6 diamino-2-phenylindole (DAPI; Sigma, St Louis, MO) to identify cell bodies. Pyramidal cells were then individually injected with Lucifer Yellow (LY; 8% in 0.1 MTris buffer, pH 7.4), and thereafter immunostained for LY using rabbit antisera against LY (1:400 000; generated at the Cajal Institute). Apical and basal dendrites were imaged at high magnification (×63 glycerol; voxel size: 0.075 × 0.075 × 0.28 *μm*^3^) using tile scan mode in a Leica TCS 4D confocal scanning laser attached to a Leitz DMIRB fluorescence microscope. Voxel size was calculated to acquire images at the highest resolution possible for the microscope (approximately 200 nanometers). Consecutive image stacks (~3) were acquired to capture the full dendritic depth, length, and width of basal dendrites, each originating from a different pyramidal neuron (10 per case; 60 dendritic segments). For apical dendrites, the main apical dendrite was scanned, at a distance of 100 *μm* from the soma up to 200 *μm* (8 dendrites per case; 16 dendritic segments). Apical and basal spines were individually reconstructed in 3D from high-resolution confocal stacks of images, using Imaris (BitplaneAG, Zurich, Switzerland), by selecting a solid surface that matched the contour of each dendritic spine. Sometimes it was necessary to use several surfaces of different intensity thresholds to capture the complete morphology of a dendritic spine, resulting in fragmented spines. Spine length was also measured, using the same software, as manual measurement from the point of insertion within the dendritic shaft to the end of the spine (see Figure 1H). Further information regarding tissue preparation, injection methodology, immunohistochemistry, imaging, 3D reconstruction, and ethics statement details is outlined in (Benavides-Piccione et al., 2013).

**Figure 1.**
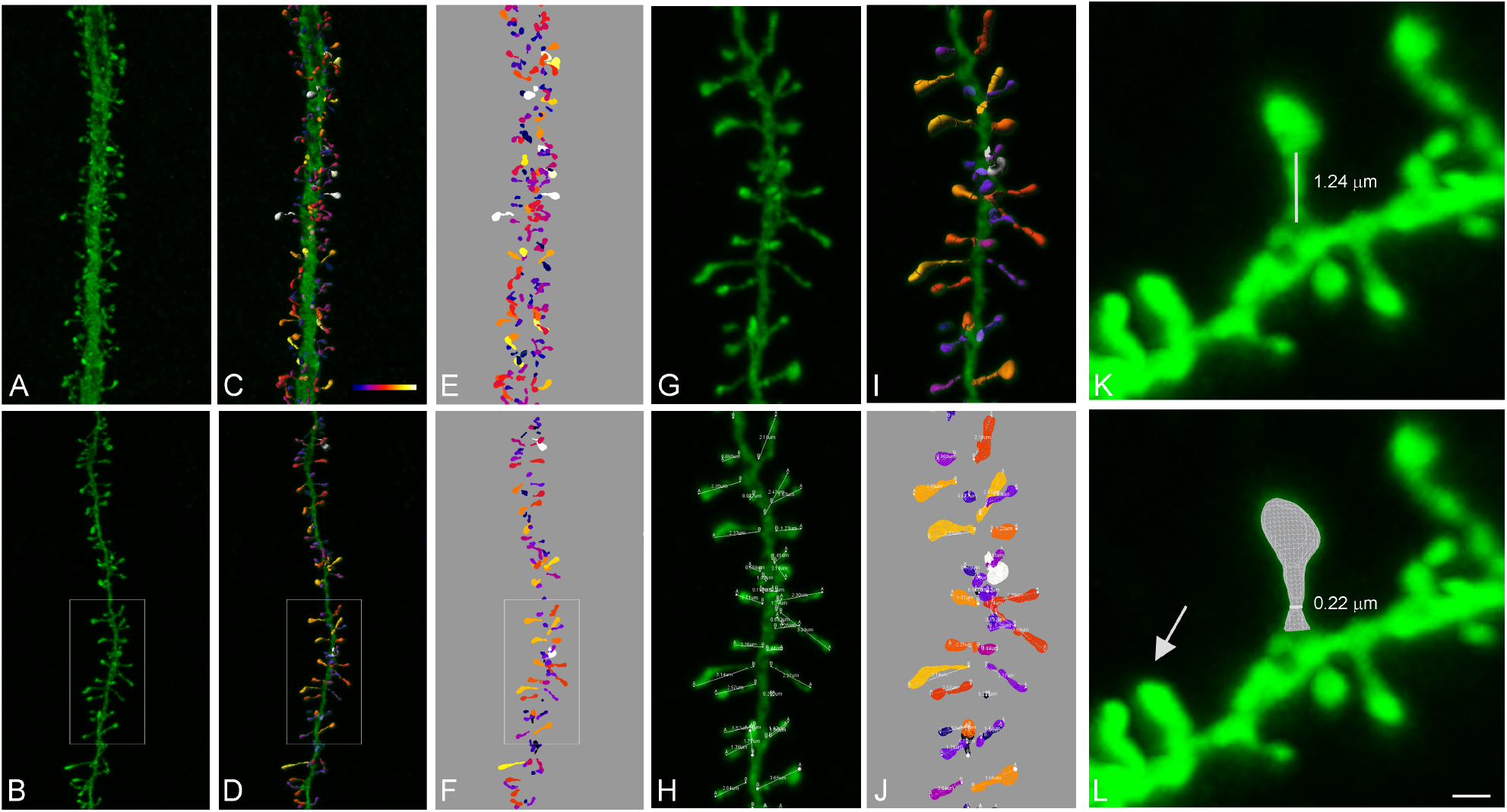
Spine reconstruction from confocal microscopy images. (A-B) Apical (A) and basal (B) dendritic segments from intracellularly injected layer 3 pyramidal neurons from the human cingulate cortex. (C-D) Shows the three-dimensional reconstruction of each dendritic spine from the dendritic segment shown in (A-B). Estimation of the spine volume values is shown by color codes (blue-white: 0-0.896 *μ*m^3^). (E-F) Shows the surface meshes that were manually created for each individual spine. (G-J) Higher magnification images of the dendritic segment shown in B. Spine lengths (that were measured in 3D) are also illustrated in H and J. (K) Example of a typical spine showing a clear head and neck. White line shows neck length. (L) Illustrates the corresponding surface mesh. White line shows neck diameter. Arrow indicates a spine not showing a clear head and neck. Dynamic scale bar (presented in L): 5 *μ*m in A-F; 2 *μ*m in G-J, and 0.6 *μ*m in K-L.

### Mouse spines dataset

Mouse tissue samples were obtained from C57BL/6 adult (8-week-old) male mice (n=4). Animals were overdosed by intraperitoneal injection of sodium pentobarbitone and perfused via the heart with phosphate-buffered saline (0.1 M) followed by 4% paraformaldehyde in PB. Brains were then removed and further immersed in 4% paraformaldehyde for 24 h. Coronal sections (200 *μm*) were obtained with a vibratome, which included the hind limb somatosensory cortical region (S1HL; Gould et al., 2012). Layer 3 pyramidal cells were then intracellularly injected and immunostained as specified above. Thereafter, apical and basal dendrites were also scanned (×63 glycerol; voxel size: 0.075 × 0.075 × 0.14 *μm*^3^; 3 apical and 7 basal dendritic segments) and a total of 1,683 spines 3D reconstructed as described above.

### Experimental design and statistical analysis

#### Spine reconstruction

The 3D mesh of each dendritic segment with all their spines was converted from the specific Imaris format (VRML file) into OBJ format files using Neuronize2 (Velasco et al., 2020). The default parameters were used: Output Resolution Percentage: 30.0, Precision: 50, Union level: 3, include segments. The reconstruct process includes joining all the pieces into one solid mesh per spine, rasterization, dilation, erosion, and a reconstruction from a binary image by 3D meshes voxelization (see Eyal et al., 2016). To correct the z-distortion caused by confocal stacks, the z-dimension values were multiplied by a factor of 0.84 (see Benavides-Piccione et al., 2013).

#### Head and neck separation

For each triangle of the mesh, we calculated two local parameters: the shape diameter function (SDF) and the distance from the triangle face to the closest point along the mesh skeleton, as described in (Ofer et al., 2021). These factors, combined with a spatial factor that takes into account the dihedral angle between neighboring faces, were applied in an energy-function graph-cut-based algorithm. Segmentation between head and neck was implemented using the Computational Geometry Algorithms Library (CGAL) 5.0.2, https://www.cgal.org “Triangulated Surface Mesh Segmentation” package. Since spines reconstructed from LM images were smoother than those reconstructed from EM, we chose the smoothing-lambda parameter to be 0.5. The morphological parameters of the head and neck were measured as described in (Ofer et al., 2021). Examples of spines separated into head and neck are presented in Figure 2. Approximately 60% of the spines could be separated into head and neck (group A in Figure 3), whereas 40% of the spines could not be clearly separated into head and neck (group B in Figure 3).

**Figure 2.**
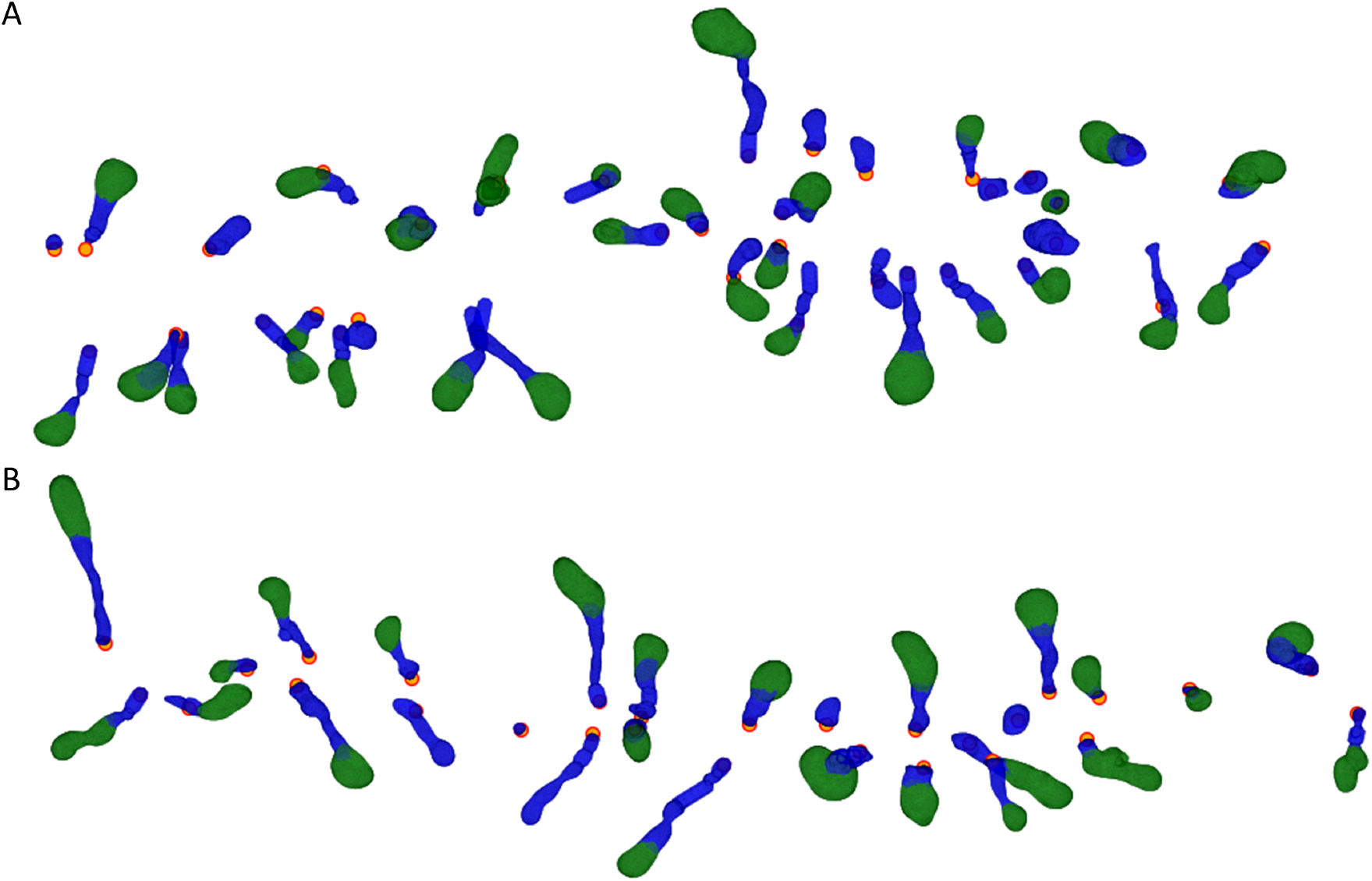
Computational separation of spines heads and necks. (A) Human apical spines, the same dendritic shaft that is presented in Figure 1A. (B) Human basal spines, the same dendritic shaft that is presented in Figure 1B. Spine heads in green and spine necks in blue. The orange dots indicate the insertion point of the spine into the dendritic shaft.

**Figure 3.**
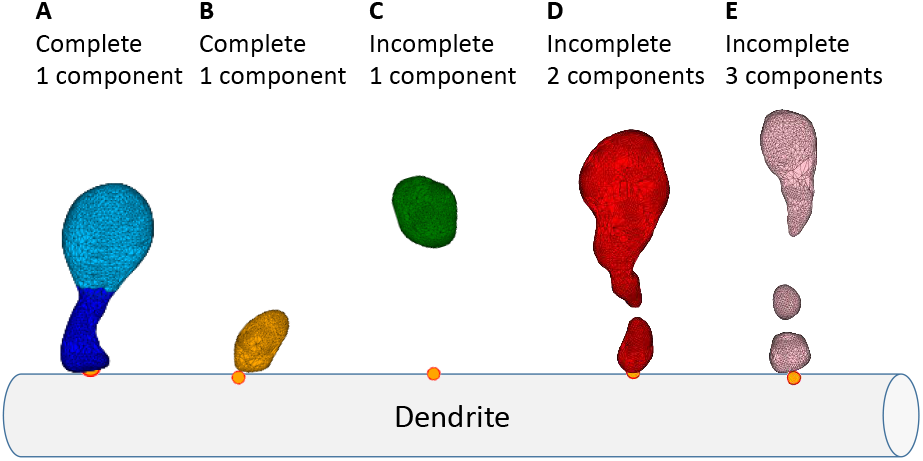
Spine dataset classification. (A) Complete spines, consisting of a single component that can be separated into the head and neck. (B) Complete spines, consisting of a single component that could not be separated into the head and neck. (C) Incomplete one component spines, detached from the dendritic shaft, containing mainly the head. (D) Incomplete spines consisting of two components. (E) Incomplete spines, consisting of three components. The orange points indicate spine insertion on the dendritic shaft.

#### Spine neck repair

A challenging process in spine morphological reconstruction is the neck, which may not be visible due to its small diameter, under the resolution limit of light microscopy. In case where it could not be detected we computationally repaired the neck. We defined five groups of spine reconstructions: (A) Complete spines attached to the dendritic shaft with a single component that can be separated into head and neck (Figure 3A); (B) Complete spines attached to the dendritic shaft, with a single component that cannot be separated into head and neck (Figure 3B); (C) Incomplete spines with one component, representing mainly the spine head, not attached to the dendritic shaft (Figure 3C); (D) Incomplete spines with two disconnected components (Figure 3D); (E) Incomplete spines with three or more disconnected components (Figure 3E).

First, we analyzed only the complete spines with clear head and neck separation (group A), representing 20% of the mouse spines and 44% of the human spines in our dataset. To further increase the number of spines in the analysis, we computationally repaired the incomplete spines (groups C and D). Adding repaired spines resulted in 60% of the spines that could be analyzed in mice and in humans. Spine meshes that contained three or more components (group E, Figure 3E) were discarded from the analysis, due to their complex shape and their small weight in the data (4% of mouse spines and 1% of human spines).

Fragmented or detached spines were reconstructed in a previous work by applying a closing morphological operator or by applying a 2D Gaussian filter (Luengo-Sanchez et al., 2018). However, since here we were interested in the separation of the spine head and neck, we developed another approach. To repair incomplete spines where the neck is invisible (group C, Figures 3C and 4A), we used the spine insertion points on the dendritic shaft (orange point in Figure 4A). When the anchor point was far away, at least 200 nm from the closest vertex of the mesh (in spines belonging to group C and not to B), we repaired the neck by adding a simple cylinder between them (Figure 4B). This enabled us to measure the neck length and allowed the separation into the head and neck. To repair disconnected spines (group D, meshes that contain two components, Figures 3D and 4D) we patched the two closest points of the separated meshes by a simple cylinder (Figure 4E), re-connecting the mesh into a single component. The radius of the cylinder used was the median neck radius measured from spines separated into head and neck before the repair process (group A), 140 nm for mice and 170 nm for humans. Although this constant radius is likely not accurate for all spines, it enabled us to segregate the head and neck, without affecting the head volume (Extended Data Figure 4-1). From repaired spines (Figure 4C and 4F; groups C and D), we measured only head volume and neck length, but not the neck diameter. Analysis of complete spines (group A) is presented in the main text of the paper, whereas analysis of both complete and repaired spines (groups A, C, and D) is presented in the Supplementary material.

**Figure 4.**
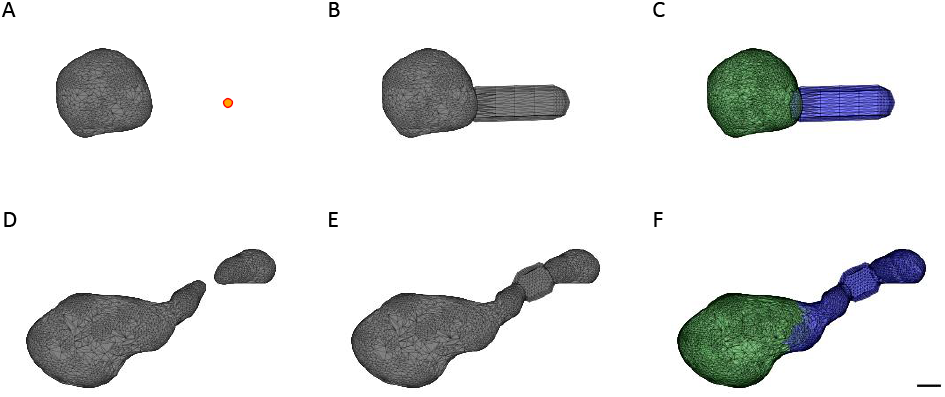
Spine neck repair of group C and group D spines. (A) Incomplete spine mesh from group C. The orange dot indicates the base point, between the spine neck and the dendritic shaft. (B) Repair of the neck by adding a cylinder between the base point and the closest vertex of the head. This cylinder is not the real neck; it was added just to enable measuring the neck length and separating the head and neck. (C) Separation into head (green) and neck (blue). (D) Incomplete spine mesh, consisting of two components (group D). (E) Repair of the neck by adding a cylinder between the two components, resulted in a connected spine. (F) Separation into head (green) and neck (blue). Scale bar: 200 nm. Extended Data Figure 4-1 shows the comparison between the morphological parameters of only the complete spines (group A) and the complete and repaired spines (groups A, C, and D).

Adding a neck (to repair group C spines) was conducted for 28% of the mouse spines and 19% of the human spines, whereas re-connection of the two-component spine (to repair group D spines) was performed in 18% of mouse spines and 12% of human spines. In some spines (5.6% in mouse spines and 3.4% in human spines) both procedures were used – a connection of the two separated components followed by a further elongation of the neck. After the repair process the human and mouse dataset contained 7,044 (89%) and 1,536 (91%) spines, respectively.

#### Statistical analysis

Mann-Whitney U rank test was used to compare groups and to calculate differences between two groups with quantitative values. Hartigan’s dip test was used to test the unimodality of the data (Hartigan and Hartigan, 1985). The correlation between parameters was examined by the Wald Test with t-distribution of the test statistic. The two-sided p-value for a hypothesis test whose null hypothesis is that the slope is zero. The asterisks indicate statistical significance: ‘n.s.’ not significant, *p< .05, ** p < .01, *** p < .001.

### Code accessibility

The CGAL scripts were written in C++; the other codes were written in Python 3.7 using the libraries numpy 1.17.4, scipy 1.5.4, and trimesh 3.9.8. Codes used are publicly available at the Columbia University Neurotechnology Center’s GitHub page (https://github.com/NTCColumbia/Spines-Morphologies-Confocal).

## RESULTS

### Human spines show dendritic and age differences in morphologies

We first compared head volume, neck length, and neck diameter in human spines samples. We found similar distributions of head volumes of basal and apical spines in the 40-years-old individual (Figure 5A, p=0.39), with a small but significant increased head volumes of basal spines of the 85-years-old individual (median of 0.356 *μm*^3^ compared with 0.333 *μm*^3^; Figure 5D, p=0.04). The neck lengths in the basal spines from 40-years-old individual were longer than those from apical spines (median of 0.607 *μm* compared with 0.566 *μm*; Figure 5B, p<.01). In the 85-years-old individual, the median neck lengths of the basal and apical spines had similar distributions (0.591 *μm* and 0.598 *μm*, respectively; Figure 5E, p=0.3). However, basal spines neck diameters were thicker than those of apical spines in samples from both the 40-years-old individual (median of 346 nm and 329 nm, respectively; Figure 5C, p<.001) and the 85-years-old individual (median of 341 nm and 321 nm, respectively; Figure 5F, p<.05).

**Figure 5.**
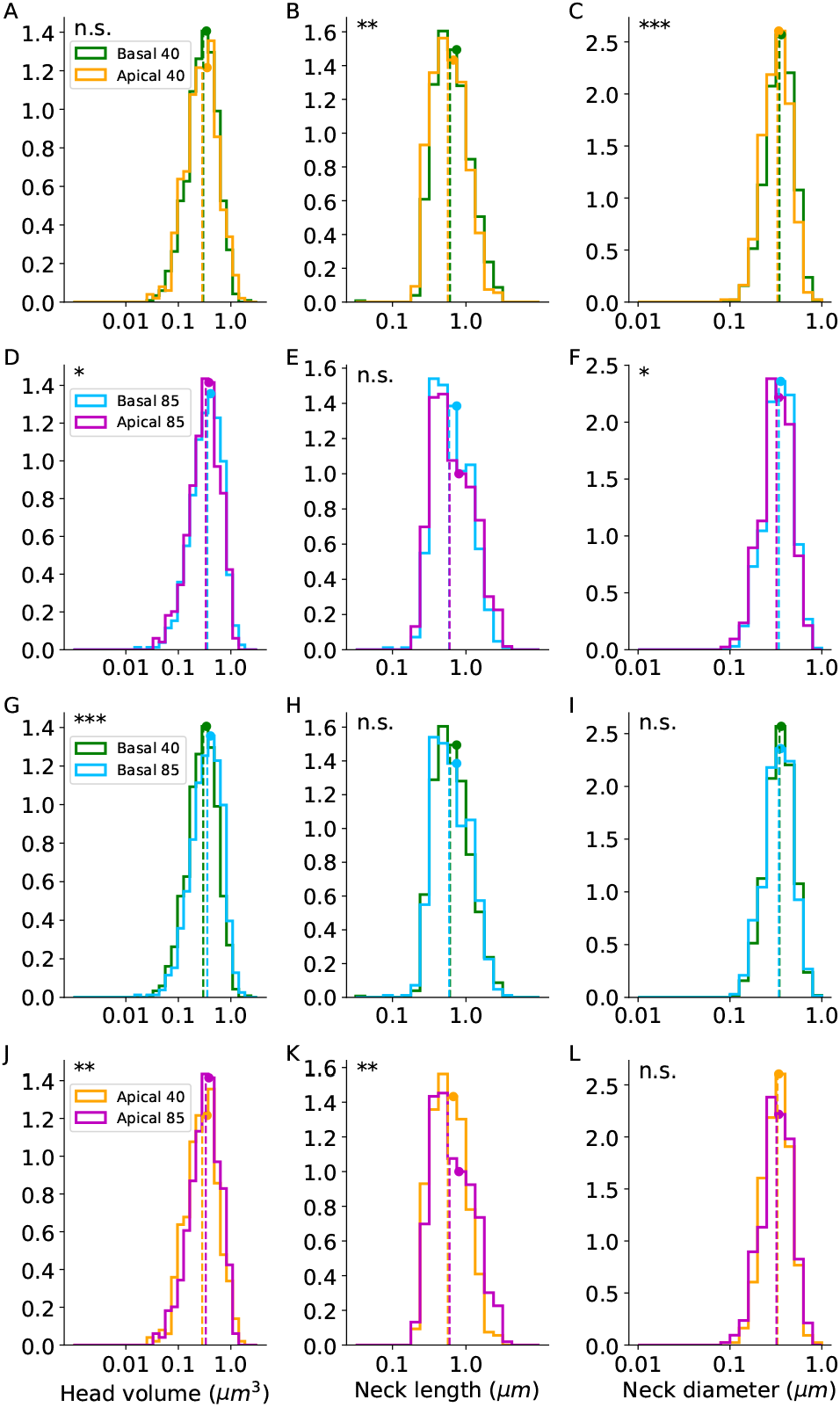
Human spines: morphological parameter distributions. The distributions of the head volumes, neck lengths, and neck diameters of complete spines (group A). (A-C) Comparison between basal and apical spines of the 40-years-old-individual. (D-F) Comparison between basal and apical spines of the 85-years-old-individual. (G-I) Comparison between 40 and 85 years-old basal spines. (J-L) Comparison between 40 and 85 years-old apical spines. Dashed vertical lines indicate the median values and the circles indicate the average. Apical 40 n=430, basal 40 n=1,012, apical 85 n=424, and basal 85 n=670. Mann-Whitney U rank test, the asterisks indicate statistical significance *p < .05, **p < .01, ***p < .001. Extended Data Figure 5-1 shows the distributions of the head volumes and neck lengths for the complete and repaired spines (groups A, C, and D).

Comparing the spines from the two individuals revealed statistically significant differences (Figure 5G-L). Basal spines of the 85-years-old individual had bigger head volumes than those from the 40-years-old individual (median of 0.356 *μm*^3^ compared with 0.298 *μm*^3^; Figure 5G, p<.001). Also, the 85-years-old apical spine heads were larger than those of the 40-years-old (median of 0.333 *μm*^3^ compared with 0.285 *μm*^3^; Figure 5J, p<.01). The neck length in basal dendrites had a similar distribution in both individuals (median of 0.591 *μm* and 0.607 *μm* in the 85 and 40 year-old individuals, respectively; Figure 5H, p=0.38), whereas in the apical spines of the 85-years-old individual, longer necks were observed (median of 0.598 *μm* compared with 0.566 *μm*; Figure 5K, p<.01). The spine neck diameters of the 85 and 40 year-old individuals were similar in basal (median of 341 nm and 346 nm, respectively; Figure 5I, p=0.2) and in apical (median of 321 nm and 329 nm, respectively; Figure 5L, p=0.49).

All the described above parameters of the complete spines (group A) including the average, standard deviation, median, and range are summarized in Table 1. The head volume and neck length values from all the spines, including also the repaired spines (groups A, C, and D), are presented in Extended Data Figure 5-1. The head volumes medians including the repaired spines were 0.298 *μm*^3^ in basal and 0.297 *μm*^3^ in apical spines of the 40-years-old individual, and 0.354 *μm*^3^ in basal and 0.341 *μm*^3^ in apical spines of the 85-years-old individual. The neck lengths medians including the repaired spines were 0.738 *μm* in basal and 0.884 *μm* in apical spines of the 40-years-old individual, and 0.734 *μm* in basal and 0.787 *μm* in apical spines of the 85-years-old individual (Table 1-1). The ranges of the parameter distributions were similar in basal and apical dendrite and for both ages.

**Table 1.** The morphological parameters values of human apical and basal spines from the 40 and 85 years-old individuals of the complete spines (group A).

| | Age | | Average ± STD | median | range |
| --- | --- | --- | --- | --- | --- |
| Head volume ($\mu m^3$) | 40 | apical | 0.3526±0.25 | 0.2846 | 0.0274 - 1.6268 |
| | | basal | 0.3426±0.22 | 0.2978 | 0.0324 - 2.206 |
| | 85 | apical | 0.3816±0.23 | 0.3331 | 0.0378 - 1.3154 |
| | | basal | 0.4103±0.25 | 0.3559 | 0.0189 - 1.538 |
| Neck length ($\mu m$) | 40 | apical | 0.6739±0.4 | 0.5664 | 0.1839 - 3.1336 |
| | | basal | 0.7464±0.46 | 0.6074 | 0.0353 - 2.9899 |
| | 85 | apical | 0.8025±0.56 | 0.5976 | 0.1865 - 3.7792 |
| | | basal | 0.7483±0.47 | 0.5911 | 0.0818 - 4.0137 |
| Neck diameter (nm) | 40 | apical | 339.9±114.36 | 328.7 | 97.4 - 836.4 |
| | | basal | 361.67±122 | 346.32 | 113.5 - 868.4 |
| | 85 | apical | 340.08±120 | 320.98 | 99.7 - 732.6 |
| | | basal | 356.66±124 | 340.62 | 100.9 - 811.8 |

In conclusion, we found differences in spine head and neck dimensions between basal and apical dendrites and between ages. Spines from the older individual had larger head volumes, in both apical and basal dendrites, and longer neck lengths (only in apical dendrites).

### Mouse spines show similar morphologies in apical and basal dendrites

We also compared the basal and apical complete spines (group A) in mice, finding similar distributions of the head volumes, neck lengths, and neck diameters (Table 2). Head volumes medians were 0.146 *μm*^3^ in basal and 0.144 *μm*^3^ in apical spines (Figure 6A, p=0.48), neck lengths medians were 0.463 *μm* in basal and 0.499 *μm* in apical spines (Figure 6B, p=0.23), and neck diameter medians were 267.1 nm in basal and 288.1 nm in apical spines (Figure 6C, p=0.2). All spines, including complete and repaired spines (groups A, C, and D), showed similar distributions of the head volumes and neck lengths of apical and basal dendrites (Extended Data Figure 6-1, Table 2-1). Head volumes medians including repaired spines were 0.124 *μm*^3^ in basal and 0.138 *μm*^3^ in apical spines, and the neck lengths medians were 0.587 *μm* in basal and 0.638 *μm* in apical. In conclusion, in mouse samples we did not find differences in spine head and neck dimensions between basal and apical dendrites, in contrast to the human data.

**Table 2.** The morphological parameters values of the apical (n=88) and basal (n=163) spines from mice of the complete spines (group A).

| | | Average ± STD | median | range |
| --- | --- | --- | --- | --- |
| Head volume ($\mu m^3$) | apical | 0.1875±0.14 | 0.1441 | 0.0151 - 0.9041 |
| | basal | 0.1832±0.13 | 0.1468 | 0.0161 - 0.7097 |
| Neck length ($\mu m$) | apical | 0.5627±0.33 | 0.4994 | 0.0195 - 2.4059 |
| | basal | 0.5467±0.31 | 0.463 | 0.2074 - 2.2687 |
| Neck diameter (nm) | apical | 295.86±116 | 288.06 | 92.41 - 640.53 |
| | basal | 278.55±102 | 267.1 | 88.61 - 676.45 |

**Figure 6.**
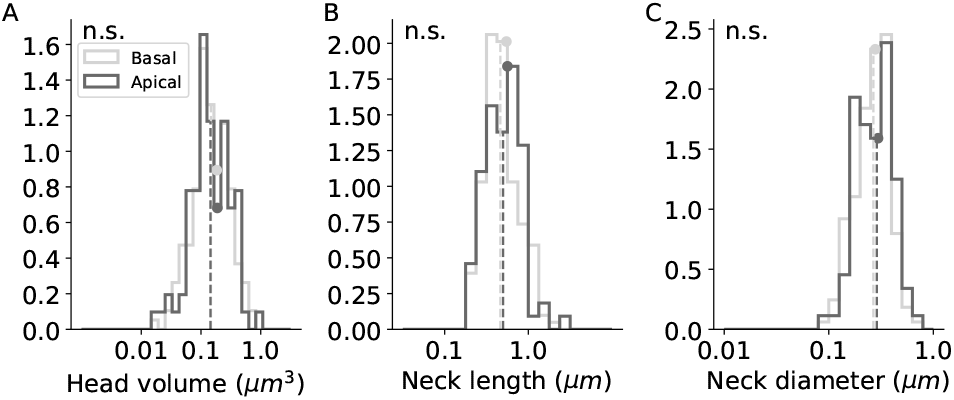
Mouse spines: morphological parameter distributions. The distributions of complete spines (group A). (A-C) Head volume, neck length, and neck diameter distributions of basal (light gray, n=163) and apical (dark gray, n=88) spines. Dashed vertical lines indicate the median values and the circles indicate the average. p=0.48, 0.23, 0.2; Mann-Whitney U rank test. Extended Data Figure 6-1 shows the results for the complete and repaired spines (groups A, C, and D).

### A continuum of spine morphologies in human and mouse spines

To explore if human spines belonged to different morphological subtypes, we analyzed the distribution of morphological features. All morphological parameters showed log-normal distributions, spreading over two or three orders of magnitude. Inspection of the distributions for each of the morphological parameters revealed skewed unimodal functions, with no clear bi-or multimodality. To test if spines could be classified into different morphological subtypes, we used Hartigan’s dip test on the head volume, neck length, and neck diameter (see Figure 7A-C; p=1, 0.99, 0.99). Then, for each pair of parameters, we explored potential multimodal distributions (Figure 7D-F), by constructing the 2-dimensional Hartigan’s dip-test that projects the scatter plot into histograms in several angles for head volume vs. neck length (p=0.83), head volume vs. neck diameter (p=0.92), and neck length vs. neck diameter (p=0.73). Further, using the 3-dimensional Hartigan’s dip-test (p=0.93), we found a continuous and unimodal distribution. The lack of multimodality was also observed when examining each population separately (basal or apical, 40 or 85 years-old individuals; not shown). Similar results, were found in the mouse dataset, with a continuum of the morphological parameters (Hartigan’s dip-test p=0.74, 0.91, 0.97; 2-dimensional Hartigan’s dip-test p=0.71, 0.44, 0.21; 3-dimensional Hartigan’s dip-test p=0.72), consistent with previous EM results (Ofer et al., 2021). Overall, the lack of multimodality in the morphological parameters is not consistent with the existence of distinct spine types. We concluded that human cortical spines, like mouse ones, displayed a continuum distribution of morphologies.

**Figure 7.**
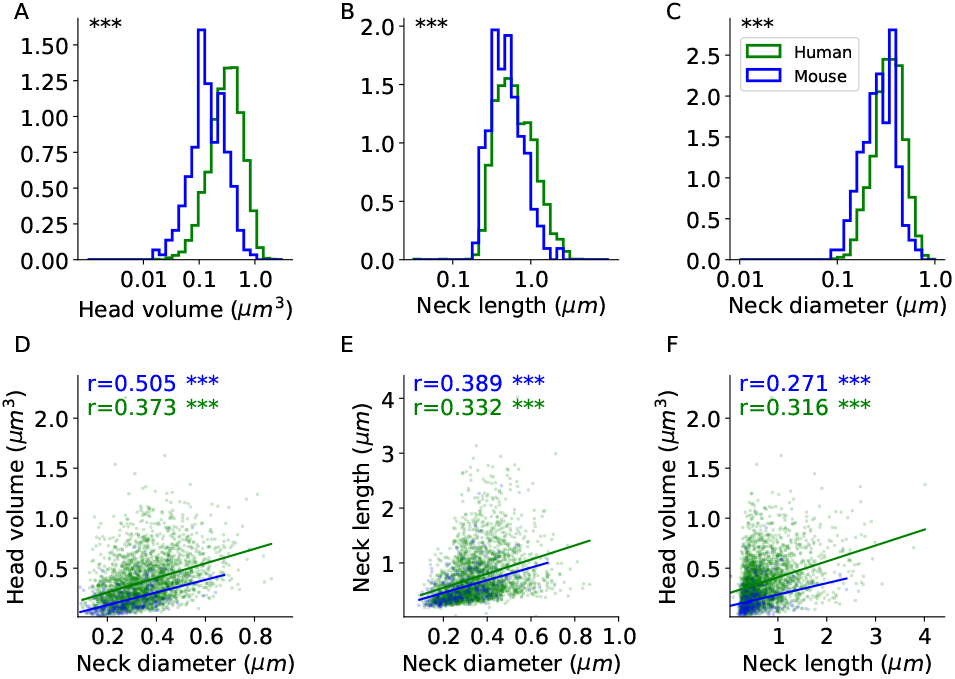
Spine morphological distributions of humans and mice. (A-C) Head volume, neck length, and neck diameter distributions in human (green, n=2,536) and mouse (blue, n=251) complete spines (group A). Mann-Whitney U rank test, ***p < .001. (D-F) Correlation between spine head and neck morphological variables. The correlation coefficients (Spearman) are indicated for each graph. The asterisks indicate statistical significance **p < .01, ***p < .001. Two-sided p-value for a hypothesis test whose null hypothesis is that the slope is zero, using Wald Test with t-distribution of the test statistic. Extended Data Figure 7-1 shows the results for the complete and repaired spines (groups A, C, and D).

### Human spines are larger and longer than those of mice

To explore species differences, we compared human and mouse complete spines (group A); (Figure 7A-C). Human spines ranged from 0.019-2.205 *μm^3^* in head volume, 0.035-4.014 *μm* in neck length, and 97.4-868 nm in neck diameter. Meanwhile, mouse spines ranged from 0.015-0.904 *μm*^3^ in head volume, 0.02-2.406 *μm* in neck length, and 89-676 nm in neck diameter. Head volumes medians were 2.19 times larger in humans than in mice (0.32 *μm*^3^ vs. 0.146 *μm*^3^), neck lengths medians were 1.27 times longer in humans (0.594 *μm* vs. 0.469 *μm*), and neck diameters medians were 1.27 times wider in humans (339 nm vs. 267 nm). When including repaired spines (groups A, C, and D), we also found bigger head volumes and longer necks in humans (Extended Data Figure 7-1A-B). We also examined the distributions of the entire spine volume and length in humans and mice. For this purpose, we also included the spines that could not be separated into head and neck (group B) with the spine that could be separated into head and neck (groups A, C, and D) (Extended Data Figure 7-1D-E). The distributions of the sphericity of the spine head were the same in mice and humans, with an average of 0.86 (Extended Data Figure 7-1F). Our results demonstrate that human spines have significantly larger head volumes and longer and thicker necks than mouse ones.

### Correlation between head and neck morphologies

Finally, we explored the existence of potential correlations between spine morphological parameters in human and mouse datasets. Consistent with previous reports (Schikorski and Stevens, 1999, 2001; Arellano et al., 2007), we found a significant positive correlation in the complete spines (group A) between head volume, neck length, and neck diameter in humans and mice (Figure 7D-F). When including also the repaired spines (groups A, C, and D), we found no correlation between head volume and neck length in mice, and only a weak correlation in humans (Extended Data Figure 7-1C). These findings are in line with previous studies in human and mouse cortex (Benavides-Piccione et al., 2002). However, in a recent study analyzing spine reconstructions from EM (Ofer et al., 2021), a weak negative correlation between neck length and neck diameter was described. Moreover, in the EM dataset there was no correlation between head volume and neck length. These differences can be explained as a result of discarding of incomplete spines (groups C and D) that present longer necks, or by the optical blurring by the LM that could enlarge the neck diameter.

## DISCUSSION

We used a computational pipeline to systematically analyze a confocal database of spines morphologies from human and mice cortical samples. These analyses supersede previous efforts, using mostly manual measurements, and enabled us to quantitatively explore potential differences in spines morphologies between apical and basal dendrites, between mice and humans, and across humans of different ages. In human samples, we find that apical spines had longer and thinner necks than basal spines, but both populations had a similar distribution of head volumes. However, no differences were observed in spine head and neck dimensions between basal and apical dendrites in mouse pyramidal neurons. Interestingly, we found that spine head volumes from older human individuals were larger, both in apical and basal compartments. There was also a significant correlation between head volume, neck length, and neck diameter in humans and mice. All morphological distributions of spine parameters, in both human and mouse samples, were unimodal, without any evidence for different spine morphological subtypes. Finally, we showed that spine head volumes, neck lengths, and neck diameters are larger in humans than in mice.

### Methodological considerations

In many samples, it was difficult to detect spine necks. Indeed, 40% of humans or mice spines could not be separated into head and neck (group B). This could be due to the fact that some spines do not have necks (stubby spines), perhaps due to the relatively young age of the mice (8 weeks) (Helm et al., 2021). But a more likely possibility is the optical blurring of head and neck morphologies, due to the resolution limit of the LM. Indeed, super-resolution stimulated emission depletion (STED) microscopy in young mice (2-5 weeks-old), shows that spines that appear stubby in LM, are in fact short-necked spines (Tønnesen et al., 2014). Consistent with this, spine morphological analyses with EM report less than 1% of spines without a clear neck (Parajuli et al., 2020; Ofer et al., 2021). In agreement with this, when examining spine volumes and lengths in all spines, including non-separated spines (group B), we find a continuum distribution (thin dashed curve in Extended Data Figure 7-1D-E), without statistical evidence for a subpopulation of spines without necks.

Comparing spine morphological parameters obtained from different microscopy methods is difficult because of methodological differences. The spine dimensions we found were different from previously reported values, especially those that used super-resolution microscopy and EM (Tønnesen et al., 2014; Levet et al., 2020). We overall measured larger head volumes, shorter neck lengths, and thicker neck diameters. These differences are expected from the lower resolution of LM, whereby a spine head may be harder for the algorithm to isolate and could include part of the neck, leading to neck shortening. Also, the tortuous structure of the spine necks may not be captured by LM meshes, and thus, be interpreted as shorter. Moreover, the blurry border of the spine confocal image could be also interpreted as an enlarged spine. However, even with these limitations, the comparisons between spine populations and between samples that we perform in this study are still valid because we used the same methodology. Finally, Table 1 correspond to a subset of the total human population of spines sampled, which include spines showing a clear differentiation between the head and neck, as well as spines in which this differentiation is not clear (Figure 2). Therefore, these values cannot be directly compared to those of other studies, where spines display a clear spine head, showing longer and thinner necks (Eyal et al., 2018).

### Differences in morphologies in human basal and apical spines

In our previous work we found differences in spines morphologies between apical and basal dendrites in human spines. Specifically, apical dendrites had higher density of spines, larger dendritic diameters, and longer spines, as compared with basal dendrites (Benavides-Piccione et al., 2013). Here, we show that apical spines have longer and thinner necks than basal spines. However, spine head volumes have similar distribution in both dendrites (see Figure 8). The similarity between basal and apical head volumes is more evident in the 40-years-old individual (Figure 5A), but also present in the 85-years-old individual, when including the repaired spines (groups A, C, and D) (Extended Data Figure 5-1A-D). The difference in neck dimension between apical and basal is also more evident in the 40-years-old (Figure 5B-C, Extended Data Figure 5-1B). The difference in neck, but not head, morphology between apical and basal spines support the argument that different mechanisms mediate the shape of the spine neck and head. Differences in spine neck dimensions could also be related to the thicker apical dendritic shaft diameter than in basal dendrites, to maintain a balance between the electrical impedance of the spine and the dendrite.

**Figure 8.**
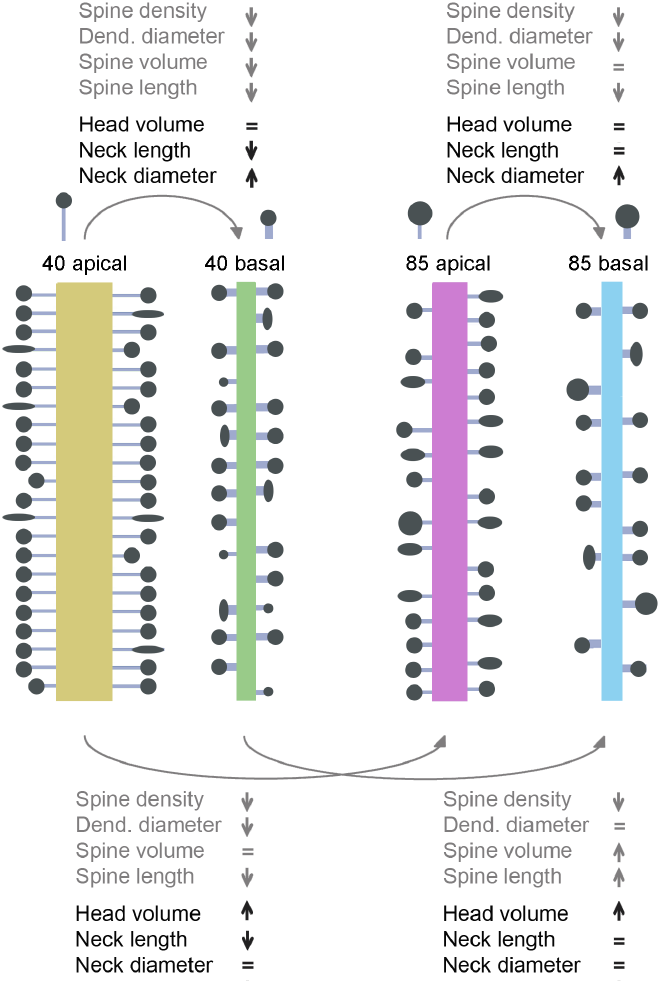
Differences in morphological parameters, including complete and repaired spines (groups A, C, and D). Apical and basal dendritic compartments of the 40-years-old and the 85-years-old human individuals. Dendrite properties from previous work (grey font; Benavides-Piccione et al., 2013) and spine properties from the present work (black font).

In contrast, in mice, both head and neck dimensions showed similar distributions in apical and basal compartments (Figure 6). Because human spines are larger than mice ones, there were fewer neck deletions (groups C and D), so the separation between head and neck was more robust and this could lead to the difference in results between humans and mice. Also, in mice, there are smaller differences in diameter between apical and basal dendritic shafts, like in hippocampus (see Figures 5A and 9A in Benavides-Piccione et al., 2020). This is consistent with the hypothesis that spine neck dimensions compensate for dendritic shaft diameters. The more similar the dendritic shaft diameters, the more similar the spine neck thickness.

### Age-related morphological differences in spines

In our previous work, we found that basal dendrites from the younger individual (40-years-old) showed more small spines, whereas in the older individual (85-years-old) basal dendrites showed larger spine volumes (Benavides-Piccione et al., 2013). Consistent with this, spines from basal dendrites of the younger subject were significantly smaller than those of the older one. In spite of these differences in basal dendrites, spines in apical dendrites were similar across age, suggesting that, with aging, small basal dendritic spines are lost, or that those spines become larger. Also, the higher percentage of small spines in basal dendrites of the 40-year-old case could be related to a higher plasticity than in the older individual. Also, in the younger individual, basal dendrites had a higher proportion of short dendritic spines, whereas apical dendrites had a higher proportion of long dendritic spines. Thus, apical and basal dendrites of the younger case showed greater differences between each other. These results suggested that shorter spines of basal dendrites and longer spines of apical dendrites are lost with age.

In the present study, we show that in the older individual, there are more spines with larger head volumes and shorter necks than in the younger individual (Extended Data Figure 5-1G-H), but the range of the distributions is similar in both individuals. This could be explained by a specific loss of ‘thin’ spines (Dumitriu et al., 2010; Dickstein et al., 2013), or by an increase in head volumes during aging. Consistent with this, changes in the distribution of the head volumes were found in aged monkeys when analyzing ‘thin’ and ‘mushroom’ spines separately (Motley et al., 2018), which implies age-related increases in spine head volume.

### A continuum of spine morphologies in human spines

We explored whether human spines belonged to morphological subtypes or are part of a continuum of morphological shapes. Statistical tests could not reject the unimodal hypothesis, meaning that we cannot prove the existence of distinct types of spines. This conclusion is of course limited to our dataset and our measured variables, so we cannot rule out the possibility that in different datasets, or with different morphological measurements, one could identify different subtypes of spines. However, the simplest interpretation of our results is that spines represent a continuum of morphologies, without any clear subtypes. This continuous distribution of spine morphologies, which we find in human and mouse samples, is consistent with our previous studies in humans using LM (Benavides-Piccione et al., 2002, 2013) and in mice using EM (Ofer et al., 2021). A recent probabilistic analysis of spine length, width, size, and curvature of 3D reconstructed human spines revealed the existence of clusters (Luengo-Sanchez et al., 2018). However, spines could not be clearly assigned to morphological classes because of the transitions between shapes. The unimodality of spine variables analyzed is inconsistent with previous attempts to classify spines into morphological subtypes, such as stubby, thin, and mushroom. These categories should instead be interpreted as shorthand description of a reality where spines have a rich distribution of different morphologies.

### Comparison of human and mouse spines

We find that human spines have bigger heads, and longer and thicker necks than mouse spines. These differences are large (Figure 7A-C), far beyond differences in spine morphologies observed between different brain regions, layers, cell types, locations along the dendritic tree, or individual ages. The morphological differences between human and mouse spines could be significant in electrical properties or information processing capabilities (Benavides-Piccione et al., 2002, 2020; Mohan et al., 2015; Fişek and Häusser, 2020; Beaulieu-Laroche et al., 2021). One can explore this with a passive electrical model, where the spine neck electrical resistance is proportional to the neck length and inversely proportional to its radius:

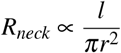

where *R_neck_* is the resistance of the neck, *l* is the neck length, and *r* is the neck radius. Using the median values of spine neck length and radius from the complete spines (group A), in humans (0.594 *μm*, 170 nm) and mice (0.469 *μm*, 134 nm), and assuming a similar cytosolic electrical resistance, we calculated the ratio between neck resistances in humans and mice of 1.27. This means that the neck resistance of mouse spines is, on average, 27% higher than that of human spines. When considering all spines (groups A, C, and D), the ratio is 1.24. Also, the ratio between apical and basal dendrites and 40-years-old and 85-years-old individuals is 1.035, 1.006, respectively, meaning that the spine necks in these samples should have similar resistances.

Although the spine necks are longer in humans than in mouse spines, the neck electrical resistance, and thus the spine head isolation, could be lower, because of thicker neck diameters. As spine neck resistance of mice spines was recently estimated to be on average 226 *M*Ω (Cornejo et al., 2022), average human spine neck resistances would be around 180 *M*Ω, based on group A spines. A previous study of human spines estimated a lower neck resistance of 50-80 *M*Ω (Eyal et al., 2018), which may be related to a different population of spines (and different spine necks diameters) included within each study.

Differences in electrical isolation of spines in humans could affect the functional of computations of individual spines. A smaller electrical isolation of human spines, together with the larger spine head, which is proportional to synaptic strength (Schikorski and Stevens, 1999, 2001; Arellano et al., 2007), implies that the functional impact of human spines, and the current that they inject into the dendrites, could be larger than those from mouse neurons (Benavides-Piccione et al., 2002), resulting in larger EPSPs in the dendrite. Human membranes also have lower capacitance and different active membrane properties that affects dendritic activity, generating larger EPSPs and reduced dendritic delay (Eyal et al., 2016; Kalmbach et al., 2018). These properties could enable more reliable synapses and the transmission of higher frequency of spike trains and support the broader range of electrical frequencies that exist in the human brain (Bragin et al., 1999; Eyal et al., 2014; Testa-Silva et al., 2014; Wang et al., 2016).

## ACKNOWLEDGMENTS

We thank Yuste Lab members for useful advice and comments. Supported by the NINDS (R01NS110422; R34NS116740) and NIMH (R01MH115900).

## Extended Data

**Figure 4-1.**
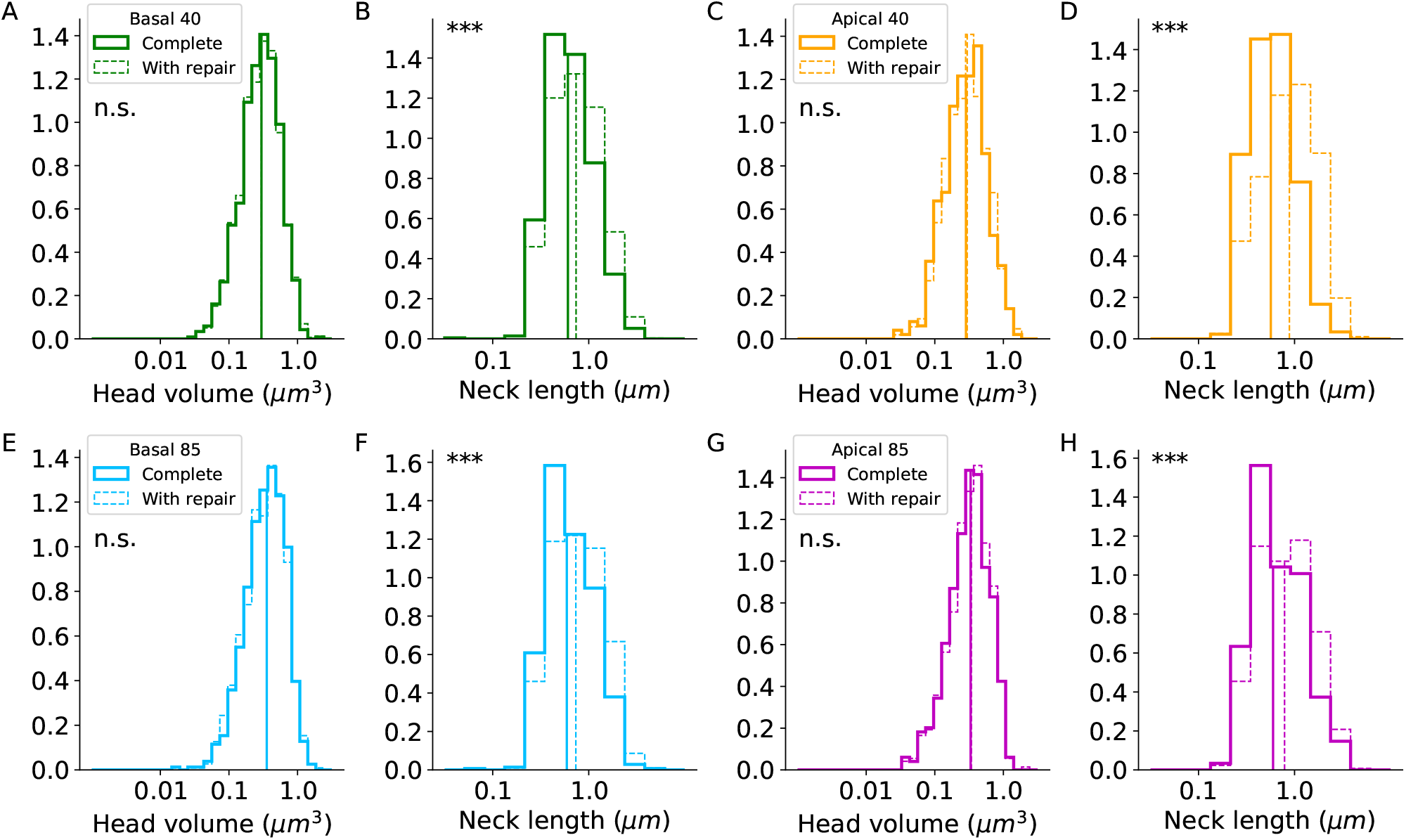
The repair process extends necks without changing head sizes. Head Volume and neck length distributions of the complete spines (group A, continuous lines) and including the repaired spines (groups A, C and D, dashed lines). (A-B) Basal spines of the 40-years-old individual, n=1,359, 1,012. (C-D) Apical spines of the 40-years-old individual, n=925, 430. (E-F) Basal spines of the 85-years-old individual, n=949, 670. (G-H) Apical spines of the 85-years-old individual, n=623, 424. Vertical lines indicate the median values. Mann-Whitney U rank test, the asterisks indicate statistical significance ***p < .001.

**Figure 5-1.**
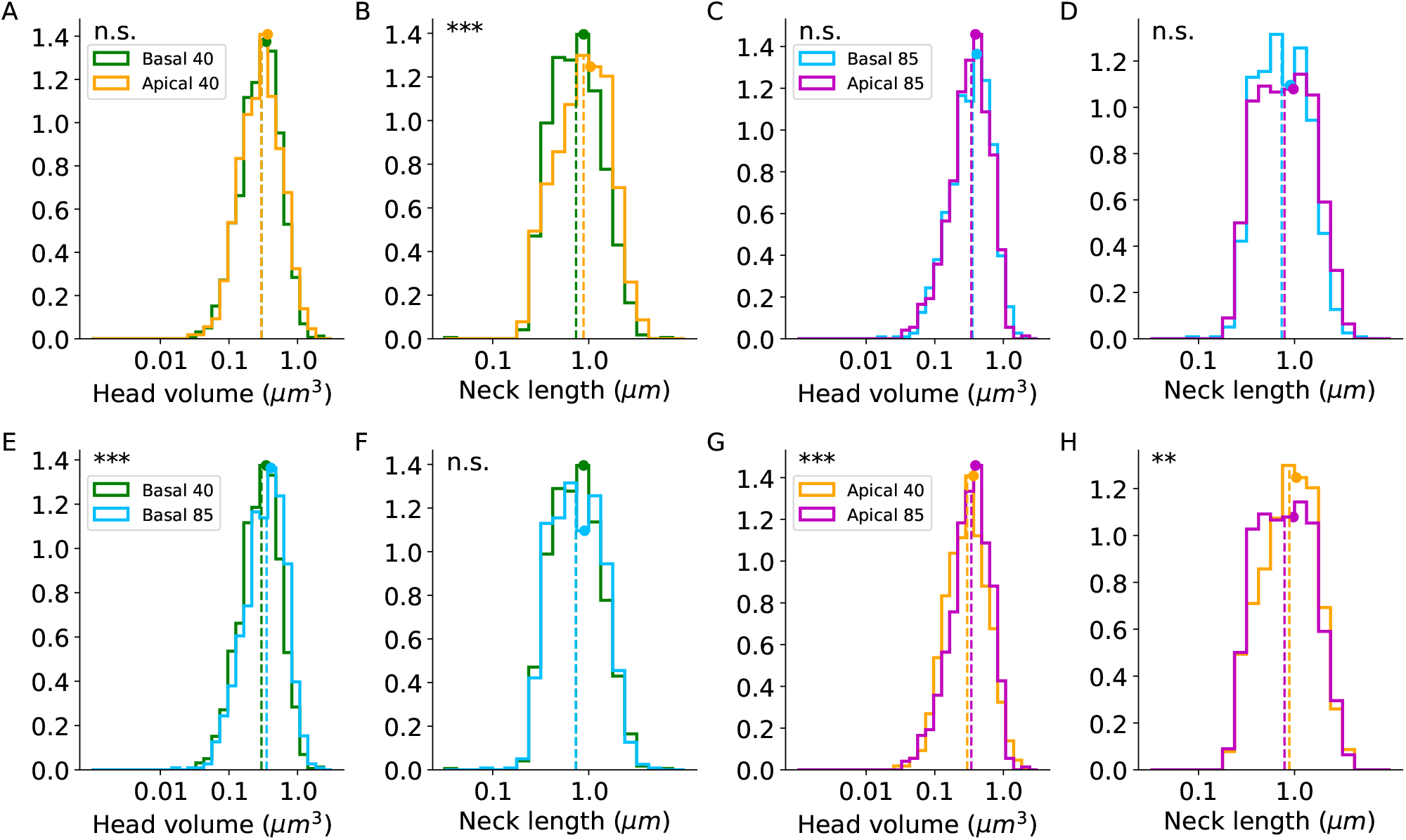
Human spines morphological parameter distributions in the complete and repaired spines (groups A, C, and D). The distributions of the head volumes and neck lengths. (A-B) Comparison between basal and apical spines of the 40-years-old-individual. (C-D) Comparison between basal and apical spines of the 85-years-old-individual. (E-F) Comparison between 40 and 85 years-old spines in the basal spines. (G-H) Comparison between 40 and 85 years-old spines in the apical spines. Dashed lines indicate the median values and the circles indicate the average. Apical 40 n=925, basal 40 n=1,359, apical 85 n=623, and basal 85 n=949. Mann-Whitney U rank test, the asterisks indicate statistical significance **p < .01, ***p < .001.

**Figure 6-1.**
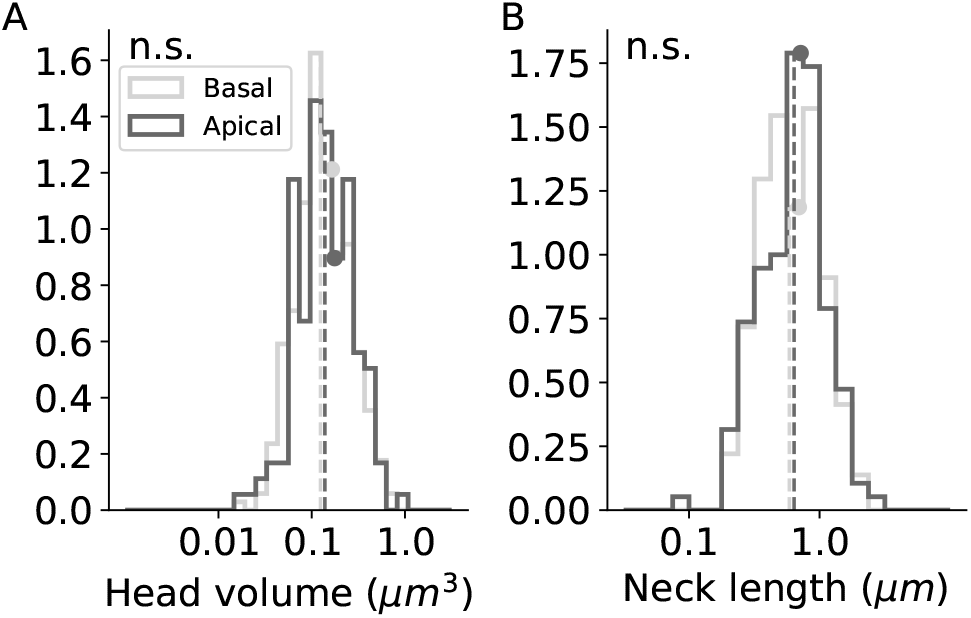
Mouse spines morphological parameter distributions in complete and repaired spines (groups A, C, and D). (A-B) Head volume and neck length distributions of basal (light gray, n=290) and apical (dark gray, n=153) dendritic spines. The parameters were calculated only for the complete spines (group A). Dashed lines indicate the median values and the circles indicate the average. p=0.17, 0.23; Mann-Whitney U rank test.

**Figure 7-1.**
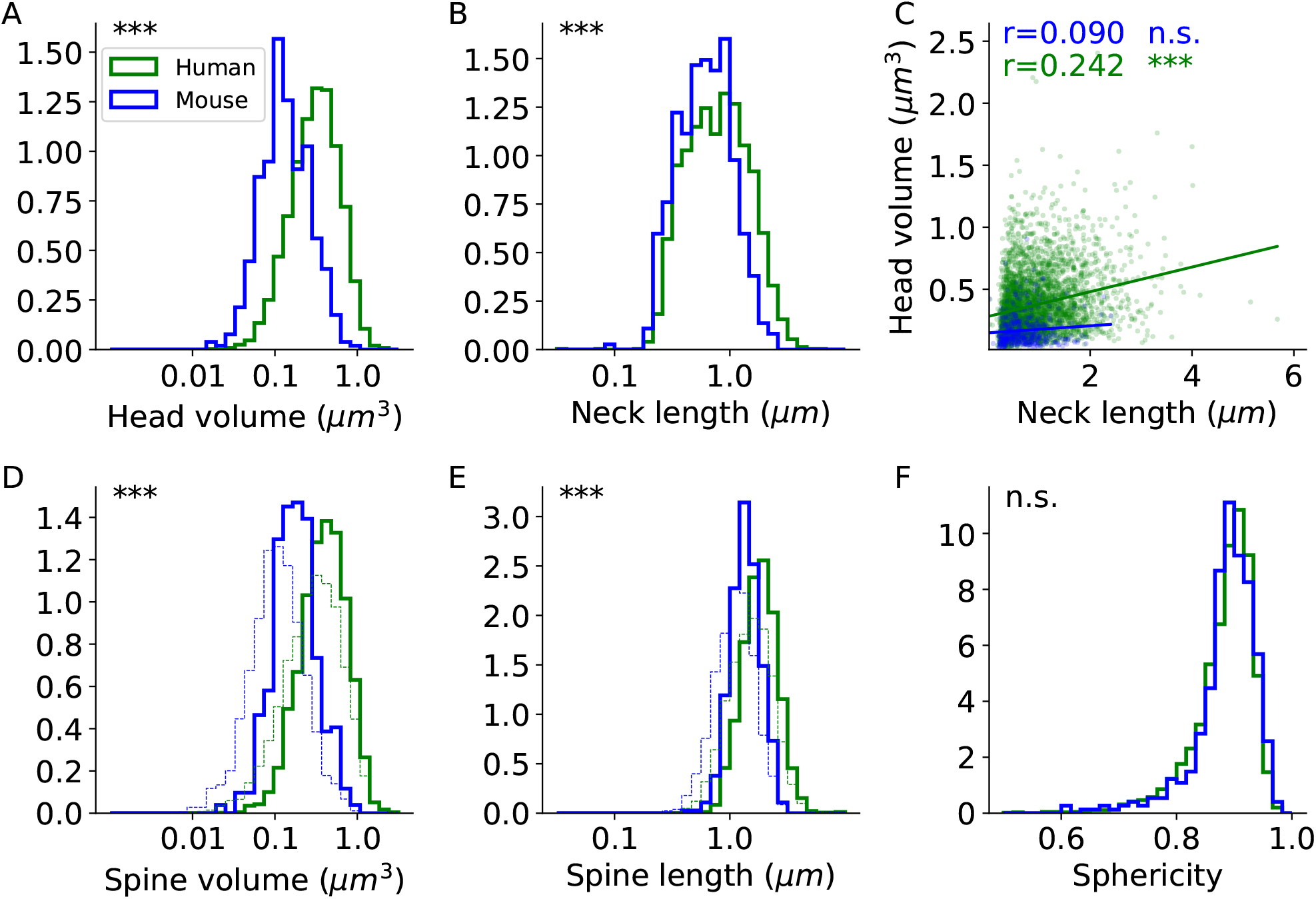
Comparing spine morphological distributions of humans and mice in complete and repaired spines. (A-B) Head volume and neck length distributions in human (green, n=3,856) and mouse (blue, n=443) complete and repaired spines (groups A, C, and D). (C) Correlation between spine head volume and neck length. The correlation coefficients (Spearman) are indicated. Two-sided p-value for a hypothesis test whose null hypothesis is that the slope is zero, using Wald Test with t-distribution of the test statistic. (D-E) Spine volume and spine length distributions in humans (n=3,856, green) and mice (n=443, blue). The thin dashed lines indicate the distribution of the all spines (groups A, B, C, and D), including those that are unseparated into head and neck, in humans (n=7,044) and mice (n=1,536). (F) The sphericity of the spine head in humans and mice (p=0.077). Mann-Whitney U rank test, the asterisks indicate statistical significance ***p < .001.

**Table 1-1.** The head volume and neck length values of the apical and basal spines from the 40 and 85 years-old individuals of the complete and repaired spines (groups A, C, and D).

| | age | | Average ± STD | median | range |
| --- | --- | --- | --- | --- | --- |
| Head volume ($\mu m^3$) | 40 | apical | 0.3665±0.26 | 0.297 | 0.0274 - 1.7596 |
| | | basal | 0.348±0.23 | 0.2977 | 0.0324 - 2.3225 |
| | 85 | apical | 0.3923±0.25 | 0.3414 | 0.0379 - 2.4035 |
| | | basal | 0.4087±0.26 | 0.3544 | 0.019 - 2.1746 |
| Neck length ($\mu m$) | 40 | apical | 1.0438±0.65 | 0.8836 | 0.0314 - 4.0011 |
| | | basal | 0.885±0.56 | 0.7381 | 0.0353 - 5.6753 |
| | 85 | apical | 0.9732±0.64 | 0.7867 | 0.1865 - 4.158 |
| | | basal | 0.9038±0.57 | 0.7342 | 0.0818 - 5.1498 |

**Table 2-1.** The head volume and neck length values of the apical (n=153) and basal (n=290) spines from mice of the complete and repaired spines (groups A, C, and D).

| | | Average ± STD | median | range |
| --- | --- | --- | --- | --- |
| Head volume ($\mu m^3$) | apical | 0.176±0.13 | 0.1384 | 0.0151 - 0.9041 |
| | basal | 0.1645±0.12 | 0.1241 | 0.0161 - 0.7097 |
| Neck length ($\mu m$) | apical | 0.7164±0.4 | 0.6382 | 0.0195 - 2.4059 |
| | basal | 0.6938±0.38 | 0.5871 | 0.2074 - 2.2687 |

## Notes

### Competing Interest Statement

The authors have declared no competing interest.

